## Supplement for "The S-component fold: A link between bacterial transporters and receptors"

#### Supplementary Information

##### Table of content

- Supplementary Figures 1-4
- Supplementary Tables 1-7
- Supplementary References

#### Supplementary Figures

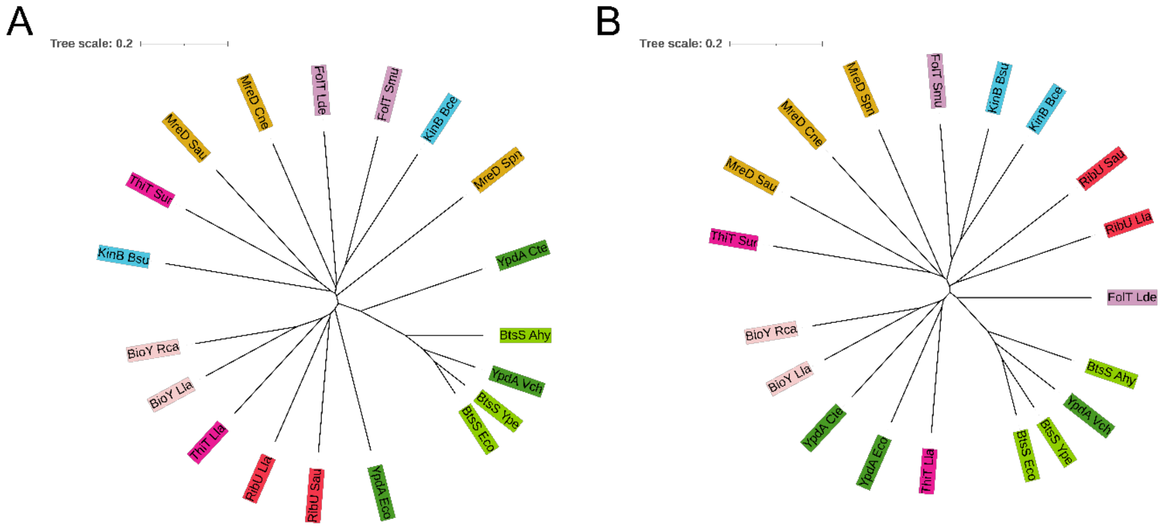


**Membrane domain of SHKs** **Full-length SHKs**

**Supplementary Figure 1** Unrooted tree of the structural multialignments of AlphaFold structures with the shared membrane six-helical fold. The AlphaFold structures of the membrane domain alone (amino acids 1-200) of sensor histidine kinases YpdA, BtsS and KinB (**A**) or their full-length version (**B**) were employed as entry to visualize the existing relationship with other SHKs, S-components (RibU, FolT, ThiT and BioY) and MreD proteins. The representative homologues come from the phyla Bacillota (Bsu = *Bacillus subtilis*, Spn = *Streptococcus pneumoniae*, Sur = *Streptococcus urinalis*, Sau = *Staphylococcus aureus*, Lde = *Lactobacillus delbrueckii*, Smu = *Streptococcus mutans*, Bce = *Bacillus cereus*, Cte = *Clostridium tetani*, Lla = *Lactococcus lactis*) and Pseudomonadota (Cne = *Cupriavidus necator*, Vch = *Vibrio cholerae*, Eco = *Escherichia coli*, Rca = *Rhodobacter capsulatus*, Ype = *Yersina pestis*, Ahy = *Aeromonas hydrophila*), and are listed more in detail on Table S5. The tree scale indicates the number of amino acid substitutions for site. The multialignments were realized via the online PROMALS3D server,[1] using standard parameters (identity threshold above which fast alignment is applied 0.6, weight for constraints derived from sequences 1.0, weight for constraints derived from homologs with structures 1.5, weight for constraints derived from input structures 1.5). Both the trees were displayed using the online Interactive Tree Of Life (iTOL) v5 tool.[2]

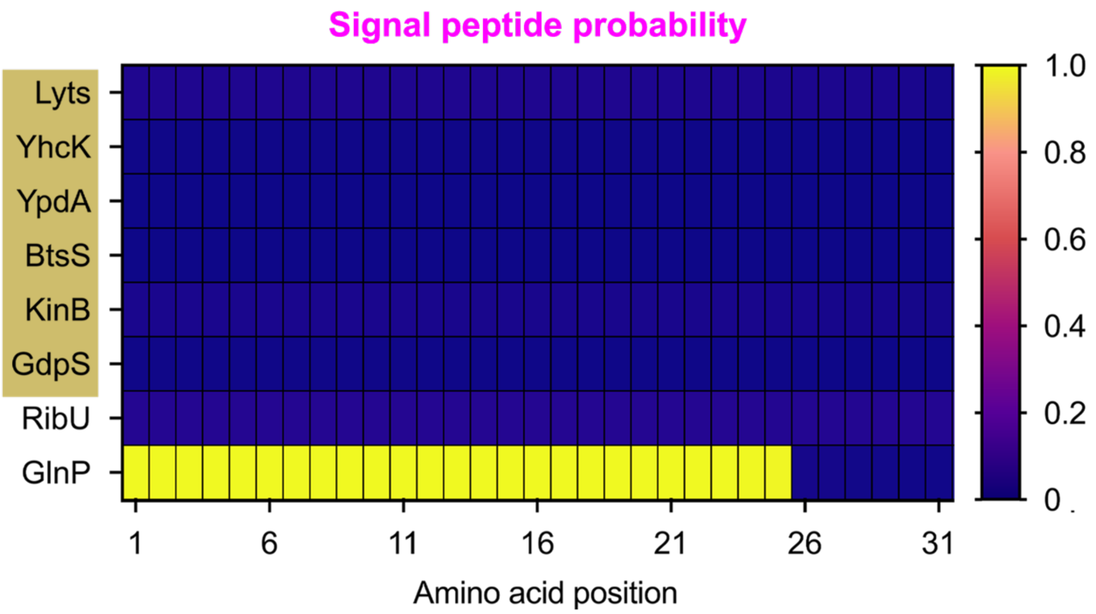


**Supplementary Figure 2** Signal peptide prediction at helix 0 of 5TMR-sensor histidine kinases. The presence of a signal peptide in the correspondence of helix 0 of LytS, YhcK, YpdA, BtsS, KinB and GdpS (UniProt accession numbers: P94513, P54595, P0AA93, P0AD14, Q08430, Q2G061) was predicted via the webserver SignalP 5.0.[3] SHKs are shaded in ochre. The sequences of RibU and GlnP (UniProt accession numbers: E5QVT2 and Q9CES5) were included in the analysis for signal peptide prediction as positive and negative controls, respectively.

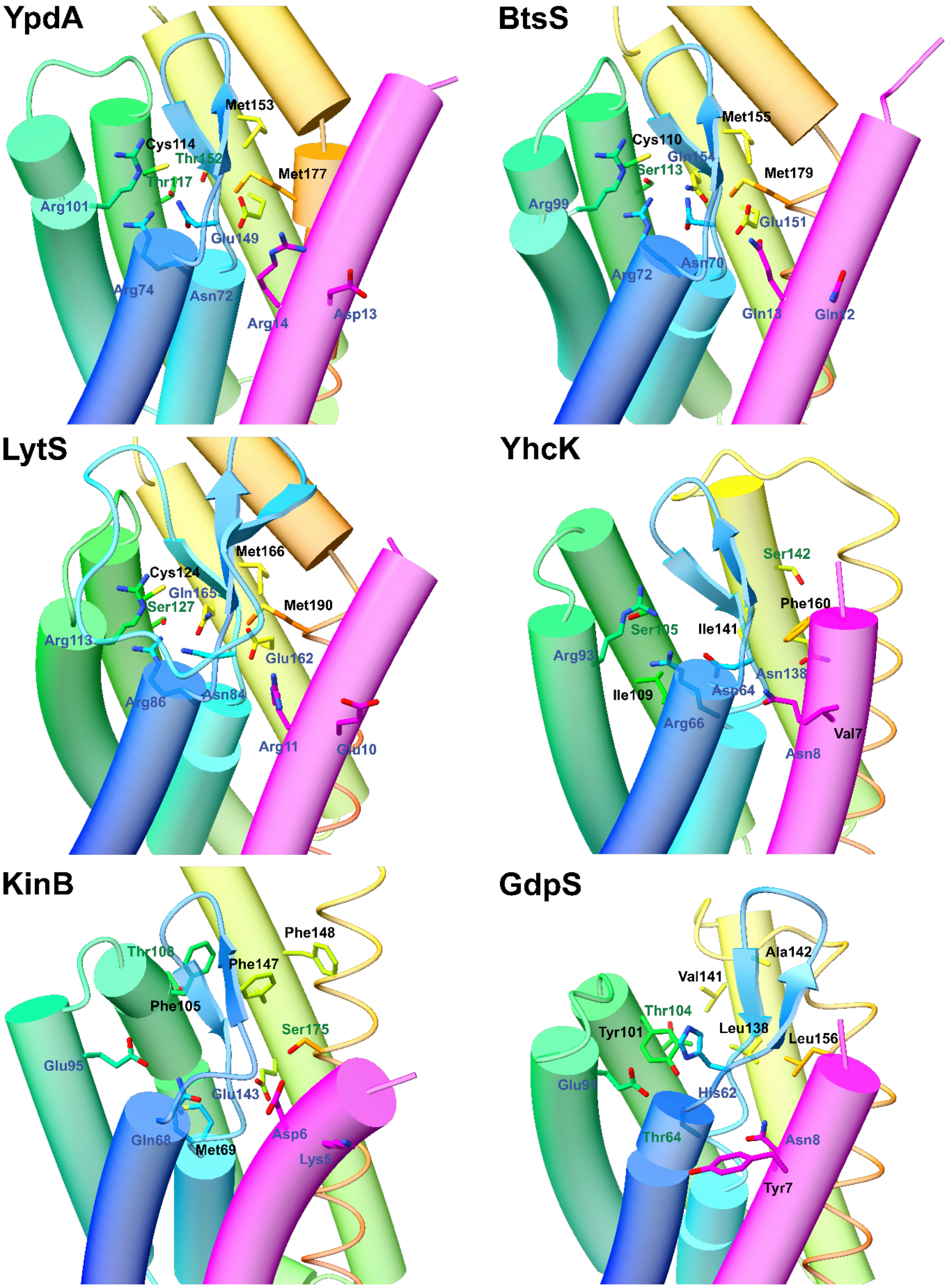


**Supplementary Figure 3** Amino acid composition of YpdA, BtsS, LytS, YhcK, KinB and GdpS corresponding to the sequence logo of 5TMR-SHKs. The additional helix 0 is highlighted in magenta, while the helices 1-6 constituting the shared fold are colored in blue, cyan, blue aqua, green, yellow and orange, respectively. The predicted AlphaFold structures, including the specific amino acid side chains, were visualized using UCSF CHIMERAX software (version 1.6.1).

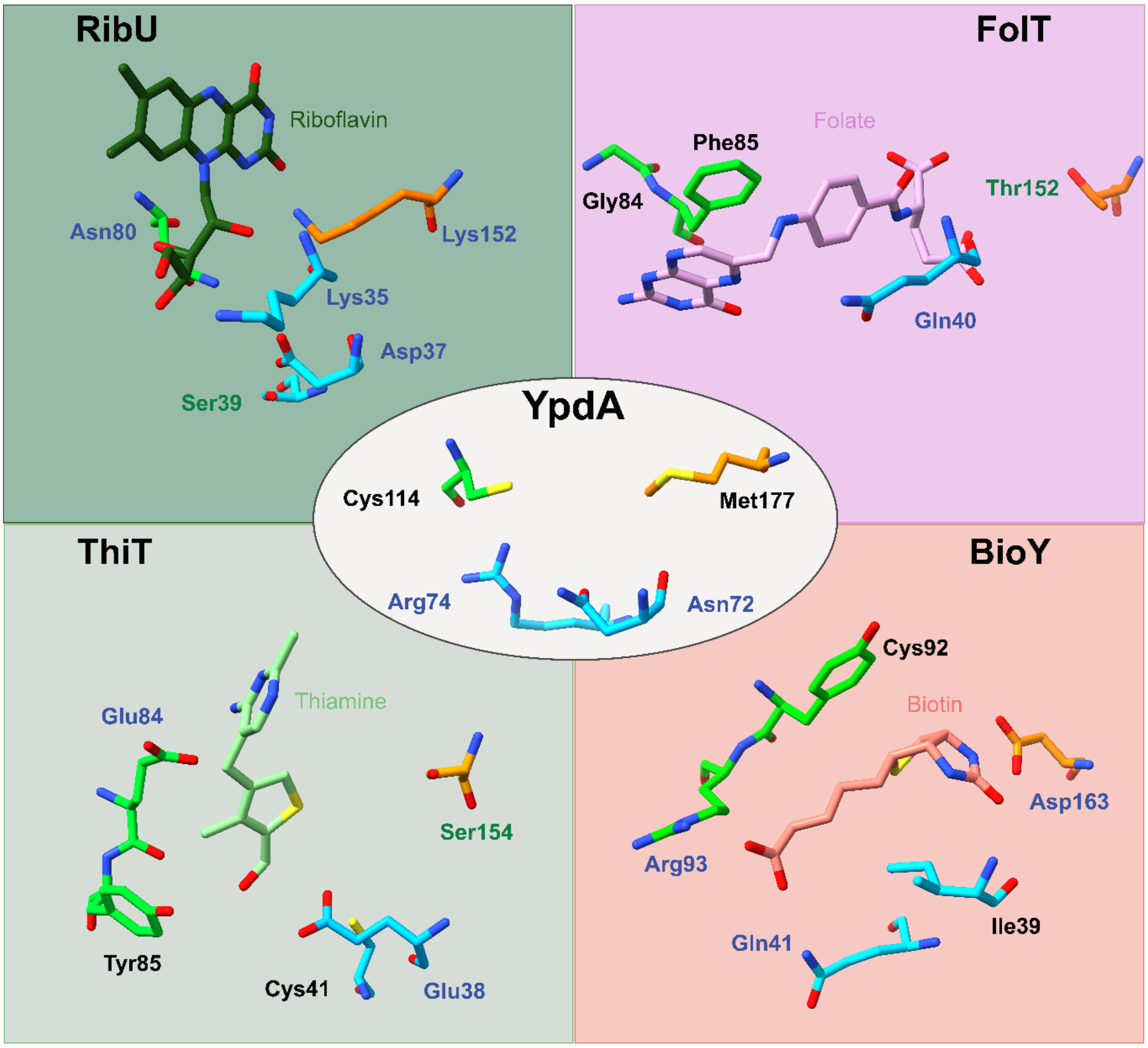


**Supplementary Figure 4** Comparison between conserved amino acids in 5TMR-SHKs and amino acids involved in substrate binding for different S-components. Using the Matchmaker tool embedded in the USCF Chimera software, the predicted structure of YpdA from *E. coli* (PDB entry: AF-P0AA93-F1) was superimposed with experimentally determined structures of S-components RibU, FolT, ThiT and BioY (PDB entries: 5KBW, 4POP, 5D0Y, 4DVE) in complex with their substrates. Asn72, Arg74, Cys114 and Met177 of the receptor YpdA resemble the tridimensional organization of particular amino acid residues in different S-components, regardless of their substrate specificity. These amino acids are located in the extracellular loop between helices 1 and 2 (in cyan), helix 4 (green) and helix 6 (in orange), and retain the same orientation towards the substrate pocket, albeit having distinct roles in substrate binding.

### Supplementary Tables

| Name | UniProt code | TCDB code | TCDB description | Substrate | Organism | Number amino acids | Reference |
|---|---|---|---|---|---|---|---|
| Gap1 | P19145 | 2.A.3.10.2 | amino acid-polyamine-organocation (apc) family | Amino acids | *Saccharomyces cerevisiae* | 602 | [4,5] |
| SNAT2 | Q9JHE5 | 2.A.18.6.4 | amino acid/auxin permease (aaap) family | Neutral amino acids | *Rattus norvegicus* | 504 | [6] |
| PATH | Q9VT04 | 2.A.18.8.3 | amino acid/auxin permease (aaap) family | Amino acids | *Drosophila melanogaster* | 471 | [7] |
| MEP2 | P41948 | 1.A.11.3.2 | ammonium transporter channel (amt) family | Ammonium | *Saccharomyces cerevisiae* | 499 | [8] |
| Ftr1 | P40088 | 2.A.108.1.1 | iron/lead transporter (ilt) family | Iron | *Saccharomyces cerevisiae* | 404 | [9] |
| Zrt1 | P32804 | 2.A.5.1.1 | zinc ($zn^{2+}$)-iron ($fe^{2+}$) permease (zip) family | Zinc | *Saccharomyces cerevisiae* | 376 | [9] |
| Pho84 | P25297 | 2.A.1.9.1 | major facilitator superfamily (mfs) | Phosphate | *Saccharomyces cerevisiae* | 587 | [10] |
| Sul1 | P38359 | 2.A.53.1.1 | sulfate permease (sulp) family | Sulphate | *Saccharomyces cerevisiae* | 859 | [11] |
| Sul2 | Q12325 | 2.A.53.1.13 | sulfate permease (sulp) family | Sulphate | *Saccharomyces cerevisiae* | 893 | [11] |
| KDELR2 | Q5ZKX9 | 9.B.191.1.8 | ER lumen protein-retaining receptor 2 | KDEL motif | *Gallus gallus* | 212 | [12] |

**Supplementary Table 1** List and main features of known transceptors.

| Query | | | Hit | | | FoldSeek results | | | | | |
|---|---|---|---|---|---|---|---|---|---|---|---|
| **Protein name** | **UniProt code** | **Structure Identifier** | **Protein name** | **PFAM entry** | **UniProt code** | **Database** | **Probability** | **Sequence Identity (%)** | **E-value** | **TM-Score** | **RMSD (Å)** |
| RibU | E5QVT2 | 3P5N | YpdA | PF07456 | P0AA93 | a | 1.00 | 17.1 | 2.42 $e^{-2}$ | 0.54 | 5.3 |
| | | | BtsS | PF07456 | P0AD14 | a | 0.94 | 11.2 | 3.32 $e^{-1}$ | 0.50 | 6.2 |
| | | | KinB | PF07456 | Q08430 | b | 0.69 | 14.6 | 9.56 $e^{-1}$ | 0.50 | 5.2 |
| | | AF-E5QVT2-F1 | YpdA | PF07456 | P0AA93 | a | 0.96 | 15.5 | 2.47 $e^{-1}$ | 0.53 | 5.8 |
| | | | BtsS | PF07456 | P0AD14 | b | 0.99 | 13.6 | 5.49 $e^{-2}$ | 0.51 | 6.7 |
| | | | KinB | PF07456 | Q08430 | b | 0.69 | 10.9 | 1.42 $e^{0}$ | 0.55 | 6.8 |
| FolT | Q1G930 | 5D0Y | YpdA | PF07456 | P0AA93 | a | 1.00 | 22.2 | 2.75 $e^{-2}$ | 0.63 | 4.1 |
| | | | BtsS | PF07456 | P0AD14 | a | 0.96 | 14.5 | 7.18 $e^{-1}$ | 0.61 | 3.8 |
| | | | KinB | PF07456 | Q08430 | b | 0.75 | 14.3 | 1.49 $e^{0}$ | 0.55 | 4.4 |
| | | AF-Q1G930-F1 | YpdA | PF07456 | P0AA93 | a | 1.00 | 19.1 | 3.62 $e^{-3}$ | 0.56 | 5.1 |
| | | | BtsS | PF07456 | P0AD14 | a | 0.98 | 13.8 | 5.77 $e^{-2}$ | 0.41 | 9.2 |
| | | | KinB | PF07456 | Q08430 | b | 0.87 | 11.3 | 5.01 $e^{-1}$ | 0.49 | 6.0 |
| ThiT | A2RI47 | 4POP | YpdA | PF07456 | P0AA93 | a | 1.00 | 16.2 | 1.16 $e^{-1}$ | 0.60 | 4.7 |
| | | | BtsS | PF07456 | P0AD14 | | -- | -- | -- | -- | -- |
| | | | KinB | PF07456 | Q08430 | b | 1.00 | 14.4 | 4.11 $e^{-2}$ | 0.55 | 5.4 |
| | | AF-A2RI47-F1 | YpdA | PF07456 | P0AA93 | b | 1.00 | 14.7 | 2.02 $e^{-1}$ | 0.63 | 4.5 |
| | | | BtsS | PF07456 | P0AD14 | | -- | -- | -- | -- | -- |
| | | | KinB | PF07456 | Q08430 | b | 1.00 | 13.9 | 1.93 $e^{-2}$ | 0.61 | 5.0 |
| BioY | A2RMJ9 | 4DVE | YpdA | PF07456 | P0AA93 | a | 0.98 | 18.3 | 3.08 $e^{-2}$ | 0.37 | 7.7 |
| | | | BtsS | PF07456 | P0AD14 | a | 0.57 | 12.5 | 2.88 $e^{-1}$ | 0.36 | 9.5 |
| | | | KinB | PF07456 | Q08430 | b | 0.94 | 15.1 | 2.00 $e^{-1}$ | 0.40 | 6.3 |
| | | AF-A2RMJ9-F1 | YpdA | PF07456 | P0AA93 | | -- | -- | -- | -- | -- |
| | | | BtsS | PF07456 | P0AD14 | a | 0.89 | 10.8 | 5.15 $e^{-1}$ | 0.54 | 5.2 |
| | | | KinB | PF07456 | Q08430 | | -- | -- | -- | -- | -- |

**Supplementary Table 2** Structural similarity results of sensor histidine kinases sharing the fold with S-components. To obtain a better insight into the structural alignments predicted by FoldSeek with SHK proteins, we looked into the predicted template modeling score (TM score) and the root mean square deviation value (RMSD value) obtained with the membrane domain of YpdA, BtsS and KinB. Both experimentally determined and AlphaFold predicted structures of RibU, FolT, ThiT and BioY were used to query multiple databases (a = AFDB-proteome, b = AFDB-SwissProt) integrated in the FoldSeek resource. The TM-score for YpdA and KinB was estimated ≥ 0.5, thus confirming the shared fold with the S-components RibU, FolT and ThiT, but not with BioY.[13] The corresponding RMSD values varied significantly, but without going beyond 7 Å in the majority of cases, indicating fold conservation of the membrane-embedded domain in view of a sequence length of 160-200 amino acid residues.[14] The predicted structure of BtsS resulted unrelated to ThiT but it was the only one to show high structural similarity with the AlphaFold structure of BioY (TM-score of 0.54 and RMSD of 5.2 Å). In the case of experimentally determined structures, the results refer to chain A. The AI-predicted structures were obtained from the version v2.3.2 of the AlphaFold database. The PFAM entry indicates the protein family to which YpdA, BtsS and KinB belong (PF07456 = 5TMR domain of 5TMR-LYT, i.e. transmembrane region of the receptors of the LytS-YhcK type). The probability of true-positive matches is comprised between 0.0 and 1.0, while sequence identity is reported in percentage. E-value describes the expected number of hits that could be found due to chance.

| | Confidence | Net Score | p-value | PairE | SolvE | Aln Score | Aln Length | Target Length | Query Length | Fold |
|---|---|---|---|---|---|---|---|---|---|---|
| **YpdA** | **CERT** | **57.035** | **9.00E-05** | **-269** | **2.3** | **248** | **160** | **164** | **200** | **4hzuS0** |
| | HIGH | 55.492 | 1.00E-04 | -488.9 | -5.4 | 139 | 187 | 430 | 200 | 1kplA0 |
| | HIGH | 55.474 | 1.00E-04 | -499.9 | -5 | 137 | 193 | 444 | 200 | 4eneA0 |
| | HIGH | 54.603 | 2.00E-04 | -424.5 | -10.1 | 127 | 192 | 504 | 200 | 3rkoC0 |
| | HIGH | 53.726 | 2.00E-04 | -444.5 | -12.1 | 108 | 180 | 612 | 200 | 3rkoB0 |
| | HIGH | 52.732 | 2.00E-04 | -503.4 | -8.8 | 101 | 181 | 605 | 200 | 4heaL0 |
| | HIGH | 50.52 | 4.00E-04 | -460.4 | -13.4 | 77 | 172 | 873 | 200 | 3zkvA0 |
| | **HIGH** | **50.415** | **4.00E-04** | **-267.2** | **0.8** | **196** | **157** | **164** | **200** | **4huqS0** |
| | **HIGH** | **50.224** | **4.00E-04** | **-251.9** | **12.9** | **255** | **160** | **168** | **200** | **3p5nA0** |
| | HIGH | 49.715 | 5.00E-04 | -433.6 | -9.9 | 91 | 195 | 467 | 200 | 4heaM0 |
| | HIGH | 48.922 | 6.00E-04 | -445.7 | -10.2 | 83 | 182 | 249 | 200 | 4fc4A0 |
| | HIGH | 48.86 | 6.00E-04 | -422.6 | -7.2 | 103 | 181 | 425 | 200 | 3nd0A0 |
| | HIGH | 48.82 | 6.00E-04 | -430.3 | -7.4 | 100 | 179 | 245 | 200 | 3d9sA0 |
| | HIGH | 48.509 | 6.00E-04 | -508 | -11.6 | 56 | 190 | 512 | 200 | 2nynA0 |
| | HIGH | 48.14 | 7.00E-04 | -511 | -16.1 | 31 | 194 | 556 | 200 | 4gmoA0 |
| | HIGH | 47.905 | 7.00E-04 | -626 | -7.4 | 40 | 192 | 279 | 200 | 5k7vA0 |
| | HIGH | 47.436 | 8.00E-04 | -581.9 | -1.5 | 79 | 173 | 395 | 200 | 6d0jA0 |
| | HIGH | 47.019 | 9.00E-04 | -500 | -14.1 | 35 | 196 | 901 | 200 | 4c0oA0 |
| | HIGH | 46.914 | 9.00E-04 | -415.4 | -7.9 | 88 | 182 | 911 | 200 | 2xwuB0 |
| | HIGH | 46.898 | 9.00E-04 | -582 | -3.7 | 66 | 167 | 197 | 200 | 5wtrA0 |
| | HIGH | 46.829 | 9.00E-04 | -340.5 | -16.4 | 68 | 178 | 483 | 200 | 6nbqB0 |
| **BtsS** | **HIGH** | **51.436** | **3.00E-04** | **-236.2** | **0.6** | **209** | **158** | **164** | **200** | **4hzuS0** |
| | HIGH | 50.035 | 4.00E-04 | -591.9 | -12.2 | 44 | 188 | 849 | 200 | 4adyA0 |
| | HIGH | 49.484 | 5.00E-04 | -454.8 | -17.6 | 52 | 191 | 690 | 200 | 1w27A0 |
| | HIGH | 49.27 | 5.00E-04 | -405.6 | -14.9 | 74 | 199 | 537 | 200 | 6en4C0 |
| | HIGH | 48.662 | 6.00E-04 | -414.7 | -7.1 | 105 | 181 | 612 | 200 | 3rkoB0 |
| | HIGH | 48.439 | 7.00E-04 | -459.9 | -5 | 100 | 185 | 504 | 200 | 3rkoC0 |
| | HIGH | 47.995 | 7.00E-04 | -520.2 | -12.1 | 48 | 197 | 288 | 200 | 6qo9A0 |
| | HIGH | 47.064 | 9.00E-04 | -482.8 | -8.6 | 70 | 176 | 368 | 200 | 6bw6A0 |
| **YhcK** | HIGH | 52.709 | 2.00E-04 | -478.9 | -13.8 | 96 | 179 | 612 | 200 | 3rkoB0 |
| | HIGH | 48.547 | 6.00E-04 | -470.1 | -9.8 | 86 | 180 | 605 | 200 | 4heaL0 |
| **LytS** | HIGH | 47.291 | 9.00E-04 | -391.4 | -9 | 101 | 181 | 612 | 200 | 3rkoB0 |
| **KinB** | **HIGH** | **53.381** | **2.00E-04** | **-503.2** | **4.5** | **175** | **164** | **170** | **200** | **5kbwA0** |
| | **HIGH** | **52.561** | **3.00E-04** | **-442.7** | **-4.1** | **147** | **161** | **164** | **200** | **4hzuS0** |
| | **HIGH** | **51.316** | **3.00E-04** | **-542.5** | **-0.8** | **126** | **165** | **189** | **200** | **4dveA0** |
| | HIGH | 51.276 | 3.00E-04 | -540.6 | -10 | 82 | 182 | 249 | 200 | 4fc4A0 |
| | HIGH | 49.104 | 6.00E-04 | -502.1 | -2.2 | 111 | 191 | 444 | 200 | 4eneA0 |
| | HIGH | 47.562 | 8.00E-04 | -517.5 | -13.5 | 46 | 184 | 309 | 200 | 3mzvA0 |
| | HIGH | 47.275 | 9.00E-04 | -458.4 | -8.3 | 84 | 179 | 612 | 200 | 3rkoB0 |
| | **HIGH** | **46.98** | **9.00E-04** | **-436** | **-8.5** | **90** | **162** | **187** | **200** | **4rfsS0** |
| **GdpS** | CERT | 61.643 | 3.00E-05 | -501 | -18.6 | 114 | 186 | 612 | 200 | 3rkoB0 |
| | CERT | 60.695 | 4.00E-05 | -492.6 | -16.9 | 118 | 185 | 504 | 200 | 3rkoC0 |
| | HIGH | 56.496 | 1.00E-04 | -539.9 | -11.8 | 100 | 189 | 605 | 200 | 4heaL0 |
| | **HIGH** | **56.235** | **1.00E-04** | **-435.6** | **-4.4** | **166** | **157** | **164** | **200** | **4hzuS0** |
| | HIGH | 53.556 | 2.00E-04 | -424.4 | -9.7 | 121 | 183 | 444 | 200 | 4eneA0 |
| | **HIGH** | **52.809** | **2.00E-04** | **-435.8** | **4.8** | **186** | **154** | **168** | **200** | **3p5nA0** |
| | HIGH | 52.707 | 2.00E-04 | -391.8 | -13.4 | 107 | 178 | 473 | 200 | 3rkoD0 |
| | HIGH | 52.696 | 2.00E-04 | -590.5 | -6.5 | 85 | 190 | 329 | 200 | 5ckrA0 |
| | HIGH | 50.993 | 4.00E-04 | -546.2 | -10.4 | 68 | 181 | 245 | 200 | 3d9sA0 |
| | HIGH | 50.085 | 4.00E-04 | -505.6 | -11.5 | 66 | 189 | 972 | 200 | 6fvbA0 |
| | HIGH | 49.939 | 5.00E-04 | -543.1 | -4 | 92 | 179 | 307 | 200 | 5jnqA0 |
| | HIGH | 49.654 | 5.00E-04 | -484.3 | -13.2 | 60 | 192 | 411 | 200 | 5wylA0 |
| | HIGH | 49.091 | 6.00E-04 | -367.6 | -14.4 | 84 | 175 | 427 | 200 | 4heaN0 |
| | HIGH | 49.013 | 6.00E-04 | -385.6 | -12.9 | 83 | 194 | 467 | 200 | 4heaM0 |
| | HIGH | 48.992 | 6.00E-04 | -680.4 | 3.1 | 84 | 170 | 194 | 200 | 5wucA0 |
| | HIGH | 48.823 | 6.00E-04 | -553.9 | -14.4 | 34 | 168 | 265 | 200 | 3ebbA0 |
| | HIGH | 47.849 | 7.00E-04 | -369.6 | -11.3 | 89 | 180 | 257 | 200 | 3tdsA0 |
| | HIGH | 47.208 | 9.00E-04 | -426.5 | -17.7 | 41 | 167 | 618 | 200 | 3c2gA0 |
| | HIGH | 47.18 | 9.00E-04 | -440.3 | -8.5 | 77 | 190 | 1041 | 200 | 3gjxA0 |
| | HIGH | 47.101 | 9.00E-04 | -405.3 | -8.2 | 91 | 166 | 249 | 200 | 4fc4A0 |

| | | | | | | | | | | |
|---|---|---|---|---|---|---|---|---|---|---|
| | HIGH | 46.995 | 9.00E-04 | -337.5 | -10.8 | 92 | 190 | 445 | 200 | 6f0kC0 |
| | HIGH | 46.981 | 9.00E-04 | -451.5 | -1.6 | 107 | 180 | 430 | 200 | 1kplA0 |
| | HIGH | 46.85 | 9.00E-04 | -517.8 | -14 | 32 | 166 | 292 | 200 | 3oyrA0 |
| | HIGH | 46.796 | 1.00E-03 | -643.9 | -8.3 | 25 | 168 | 279 | 200 | 5k7vA0 |
| | HIGH | 46.775 | 1.00E-03 | -460.7 | -14.1 | 43 | 185 | 1017 | 200 | 4hatC0 |
| | HIGH | 46.768 | 1.00E-03 | -511.7 | -7.2 | 62 | 186 | 836 | 200 | 4fddA0 |
| | HIGH | 46.689 | 1.00E-03 | -332.2 | -9.2 | 101 | 178 | 1044 | 200 | 4dx5A0 |

**Supplementary Table 3** pGenTHREADER (Profile Based Fold Recognition) results from the primary structures of YpdA, BtsS, LytS, YhcK, KinB and GdpS. Experimentally determined structures of S-components are highlighted in red. PairE shows the pairwise energy, SolvE means solvation energy, Aln is the abbreviation for the word 'alignment'.

| | Confidence | Net Score | p-value | PairE | SolvE | Aln Score | Aln Length | Target Length | Query Length | Region Start | Region End | CATH |
|---|---|---|---|---|---|---|---|---|---|---|---|---|
| **YpdA** | **HIGH** | **6.053** | **1.00E-05** | **-249.3** | **15.9** | **166** | **157** | **168** | **200** | **41** | **200** | **3p5nA00** |
| | **HIGH** | **5.855** | **1.00E-05** | **-299.7** | **14.1** | **161** | **158** | **171** | **200** | **41** | **199** | **5kbwB00** |
| | **HIGH** | **5.651** | **2.00E-05** | **-276.5** | **13.6** | **156** | **161** | **164** | **200** | **37** | **197** | **4hzuS00** |
| | **HIGH** | **5.401** | **3.00E-05** | **-283.3** | **11.5** | **150** | **153** | **187** | **200** | **40** | **200** | **4rfsS00** |
| | **HIGH** | **4.64** | **6.00E-05** | **-251** | **17.5** | **133** | **147** | **155** | **200** | **43** | **194** | **5d0yA00** |
| | HIGH | 4.511 | 7.00E-05 | -310.9 | 10.2 | 130 | 161 | 183 | 200 | 40 | 200 | 4tkrA00 |
| **BtsS** | HIGH | 5.191 | 3.00E-05 | -285.1 | 3.9 | 145 | 172 | 430 | 200 | 1 | 200 | 1kplA00 |
| | **HIGH** | **4.692** | **6.00E-05** | **-194.2** | **12.8** | **134** | **164** | **187** | **200** | **37** | **200** | **4rfsS00** |
| | **HIGH** | **4.465** | **7.00E-05** | **-241.2** | **9.5** | **129** | **156** | **171** | **200** | **39** | **200** | **5kbwB00** |
| | **HIGH** | **4.322** | **9.00E-05** | **-214.6** | **11.9** | **126** | **160** | **164** | **200** | **34** | **200** | **4hzuS00** |
| | **HIGH** | **4.313** | **9.00E-05** | **-160.9** | **18.6** | **126** | **160** | **168** | **200** | **39** | **200** | **3p5nA00** |
| **YhcK** | **HIGH** | **4.959** | **4.00E-05** | **-370** | **15.8** | **140** | **156** | **168** | **200** | **33** | **200** | **3p5nA00** |
| **LytS** | **HIGH** | **4.96** | **4.00E-05** | **-190.3** | **15.6** | **140** | **149** | **168** | **200** | **37** | **200** | **3p5nA00** |
| | HIGH | 4.42 | 8.00E-05 | -294 | 7.5 | 128 | 176 | 430 | 200 | 1 | 200 | 1kplA00 |
| | **HIGH** | **4.277** | **9.00E-05** | **-216.8** | **9.7** | **125** | **150** | **171** | **200** | **37** | **200** | **5kbwB00** |
| **KinB** | **HIGH** | **5.78** | **2.00E-05** | **-522.8** | **8.4** | **159** | **165** | **171** | **200** | **34** | **198** | **5kbwB00** |
| | **HIGH** | **5.522** | **2.00E-05** | **-398.7** | **16.7** | **153** | **165** | **168** | **200** | **34** | **200** | **3p5nA00** |
| **GdpS** | **HIGH** | **5.574** | **2.00E-05** | **-446.9** | **8.2** | **154** | **155** | **168** | **200** | **29** | **183** | **3p5nA00** |
| | **HIGH** | **5.274** | **3.00E-05** | **-443.9** | **9.3** | **147** | **155** | **171** | **200** | **29** | **183** | **5kbwB00** |
| | **HIGH** | **4.878** | **5.00E-05** | **-435.6** | **8** | **138** | **158** | **164** | **200** | **26** | **191** | **4hzuS00** |

**Supplementary Table 4** pDomTHREADER (Protein Domain Fold Recognition) results from the primary structures of YpdA, BtsS, LytS, YhcK, KinB and GdpS. Experimentally determined structures of S-components are highlighted in red. CATH refers to the entry of the CATH Protein Structure Classification database describing the hit.

| PFAM entry | Protein | UniProt | Structure | Organism | Phylum | Gram |
|---|---|---|---|---|---|---|
| PF12822 | RibU | E5QVT2 | AF-E5QVT2-F1 | *Staphylococcus aureus* | Bacillota | + |
| PF12822 | RibU | P0CI36 | AF-P0CI36-F1 | *Lactococcus lactis* | Bacillota | + |
| PF12822 | FolT | Q1G930 | AF-Q1G930-F1 | *Lactobacillus delbrueckii* | Bacillota | + |
| PF12822 | FolT | Q8DV98 | AF-Q8DV98-F1 | *Streptococcus mutans* | Bacillota | + |
| PF12822 | ThiT | A2RI47 | AF-A2RI47-f1 | *Lactococcus lactis* | Bacillota | + |
| PF12822 | ThiT | G5KHI3 | AF-G5KHI3-F1 | *Streptococcus urinalis* | Bacillota | + |
| PF12822 | BioY | A2RMJ9 | AF-A2RMJ9-F1 | *Lactococcus lactis* | Bacillota | + |
| PF12822 | BioY | D5ARG8 | AF-D5ARG8-F1 | *Rhodobacter capsulatus* | Pseudomonadota | - |
| PF07694 | BtsS | P0AD14 | AF- P0AD14 -F1 | *Escherichia coli* | Pseudomonadota | - |
| PF07694 | BtsS | A0KLH6 | AF- A0KLH6-F1 | *Aeromonas hydrophila* | Pseudomonadota | - |
| PF07694 | BtsS | A0A0E1NT14 | AF- A0A0E1NT14-F1 | *Yersinia Pestis* | Pseudomonadota | - |
| PF07694 | YpdA | P0AA93 | AF-P0AA93-F1 | *Escherichia coli* | Pseudomonadota | - |
| PF07694 | YpdA | Q892V8 | AF-Q892V8-F1 | *Clostridium tetani* | Bacillota | + |
| PF07694 | YpdA | Q9KU35 | AF-Q9KU35-F1 | *Vibrio cholerae* | Pseudomonadota | - |
| PF07694 | KinB | Q08430 | AF-Q08430-F1 | *Bacillus subtilis* | Bacillota | + |
| PF07694 | KinB | A0A7D4D8N3 | AF-A0A7D4D8N3-F1 | *Bacillus cereus* | Bacillota | + |
| PF04093 | MreD | Q2FXS7 | AF-Q2FXS7-F1 | *Staphylococcus aureus* | Bacillota | + |
| PF04093 | MreD | Q8DMY3 | AF-Q8DMY3-F1 | *Streptococcus pneumoniae* | Bacillota | + |
| PF04093 | MreD | Q0KFF4 | AF-Q0KFF4-F1 | *Cupriavidus necator* | Pseudomonadota | - |

**Supplementary Table 5** Identifiers and taxonomic origin of protein homologues with the shared S-component fold used for structural alignment.

| UniProt | Protein | Domain | PFAM entry | AlphaFold Structure | Organism | Phylum | Gram |
|---|---|---|---|---|---|---|---|
| Q2JYM9 | Hypothetical conserved protein | 5TM-5TMR_LYT<br>PAS_4<br>PAS_7<br>GGDEF | PF07694<br>PF08448<br>PF12860<br>PF00990 | AF-Q2JYM9-F1 | *Rhizobium etli* | Pseudomonadota | - |
| Q488U0 | Sensory box/GGDEF/EAL domain protein | 5TM-5TMR_LYT<br>PAS_4<br>GGDEF<br>EAL | PF07694<br>PF08448<br>PF00990<br>PF00563 | AF- Q488U0-F1 | *Colwellia psychrerythraea* | Pseudomonadota | - |
| F5RD42 | Sensory box protein/GGDEF domain protein | 5TM-5TMR_LYT<br>PAS_4<br>PAS_9<br>GGDEF<br>EAL | PF07694<br>PF08448<br>PF13426<br>PF00990<br>PF00563 | AF- F5RD42-F1 | *Methyloversatilis universalis* | Pseudomonadota | - |
| P54595 | Uncharacterized protein YhcK | 5TM-5TMR_LYT<br>GGDEF | PF07694<br>PF00990 | AF- P54595-F1 | *Bacillus subtilis* | Bacillota | + |
| C6WYA8 | Diguanylate cyclase/phosphodiesterase | 5TM-5TMR_LYT<br>GGDEF<br>EAL | PF07694<br>PF00990<br>PF00563 | AF- C6WYA8-F1 | *Methylotenera mobilis* | Pseudomonadota | - |
| Q64AC4 | PPM-type phosphatase domain-containing protein | 5TM-5TMR_LYT<br>SpoIIE<br>(PPM phosphatase) | PF07694<br>PF07728 | AF- Q64AC4-F1 | uncultured *archaeon GZfos32E7* | Archaea (domain) | |
| E8TNX1 | Histidine kinase | 5TM-5TMR_LYT<br>PAS_4<br>HK<br>HK-ATPase<br>REC<br>REC | PF07694<br>PF08448<br>PF00512<br>PF02518<br>PF00072<br>PF00072 | AF- E8TNX1-F1 | *Mesorhizobium ciceri biovar biserrulae* | Pseudomonadota | - |
| A8TXH3 | Histidine kinase | 5TM-5TMR_LYT<br>HK<br>HK-ATPase<br>REC | PF07694<br>PF00512<br>PF02518<br>PF00072 | AF- A8TXH3-F1 | *alpha proteobacterium BAL199* | Pseudomonadota | - |
| Q08430 | Sporulation kinase B KinB | 5TM-5TMR_LYT<br>HK<br>HK-ATPase | PF07694<br>PF00512<br>PF02518 | AF- Q08430-F1 | *Bacillus subtilis* | Bacillota | + |
| A0A4R3E8R1 | Methyl-accepting chemotaxis sensory transducer with Pas/Pac sensor | 5TM-5TMR_LYT<br>PAS_3<br>MCP | PF07694<br>PF08447<br>PF00015 | AF- A0A4R3E8R1-F1 | *Bosea species* | Pseudomonadota | - |
| A2SU24 | Signal transduction histidine kinase, LytS | NRAMP-1<br>5TM-5TMR_LYT | PF01566<br>PF07694 | AF- A2SU24-F1 | *Methanocorpusculum labreanum* | Archaea (domain) | |
| P0AA93 | Sensor histidine kinase YpdA | 5TM-5TMR_LYT<br>GAF<br>HK<br>HK-ATPase | PF07694<br>PF01590<br>PF06580<br>PF02518 | AF- P0AA93-F1 | *Escherichia coli* | Pseudomonadota | - |

**Supplementary Table 6** Identifiers, features and taxonomic origin of representative 5TMR-sensor histidine kinases with the shared S-component fold and different intracellular domain(s).

| UniProt Code | Name | Species | Length (AAs) |
|---|---|---|---|
| A0A085FHL7 | Methyl-accepting transducer domain-containing protein | *Bosea sp. LC85* | 610 |
| A0A0K6HHJ5 | Methyl-accepting chemotaxis sensory transducer with Pas/Pac sensor | *Chelatococcus sambhunathii* | 620 |
| A0A0N0MAL0 | Methyl-accepting transducer domain-containing protein | *Bosea vaviloviae* | 621 |
| A0A0Q3T337 | Methyl-accepting transducer domain-containing protein | *Bosea thiooxidans* | 620 |
| A0A0Q9IX50 | Methyl-accepting transducer domain-containing protein | *Bosea sp. Root483D1* | 608 |
| A0A126P016 | Methyl-accepting transducer domain-containing protein | *Bosea sp. PAMC 26642* | 612 |
| A0A1B3NPR9 | Sensory box protein | *Bosea sp. RAC05* | 611 |
| A0A1D7TYC1 | Methyl-accepting transducer domain-containing protein | *Bosea vaviloviae* | 611 |
| A0A1E1V000 | Methyl-accepting transducer domain-containing protein | *Bosea sp. BIWAKO-01* | 610 |
| A0A1I1PIF5 | Methyl-accepting transducer domain-containing protein | *Bosea sp. CRIB-10* | 611 |
| A0A1I3PID7 | Methyl-accepting chemotaxis sensory transducer with Pas/Pac sensor | *Bosea sp. OK403* | 611 |
| A0A1N7BQL6 | Methyl-accepting transducer domain-containing protein | *Bosea sp. TND4EK4* | 610 |
| A0A258A033 | Methyl-accepting transducer domain-containing protein | *Bosea sp. 12-68-7* | 642 |
| A0A2S4LV10 | Methyl-accepting chemotaxis sensory transducer with Pas/Pac sensor | *Bosea psychrotolerans* | 611 |
| A0A2T1HX49 | Methyl-accepting transducer domain-containing protein | *Alsobacter soli* | 610 |
| A0A2T4YBA7 | Methyl-accepting transducer domain-containing protein | *Bosea sp. 124* | 637 |
| A0A2T4YBB9 | Methyl-accepting chemotaxis sensory transducer with Pas/Pac sensor | *Bosea sp. 124* | 611 |
| A0A2W4SRZ7 | Methyl-accepting transducer domain-containing protein | *Hyphomicrobiales bacterium* | 611 |
| A0A2W6YRT7 | Methyl-accepting transducer domain-containing protein | *Chelatococcus sp.* | 610 |
| A0A370L925 | Methyl-accepting transducer domain-containing protein | *Bosea caraganae* | 998 |
| A0A4Q1TE83 | Methyl-accepting transducer domain-containing protein | *Bosea sp. Tri-39* | 629 |
| A0A4Q1UWR4 | Methyl-accepting transducer domain-containing protein | *Bosea sp. Tri-44* | 629 |
| A0A4Q3FEP4 | Methyl-accepting transducer domain-containing protein | *Hyphomicrobiales bacterium* | 638 |
| A0A4R3E8R1 | Methyl-accepting chemotaxis sensory transducer with Pas/Pac sensor | *Bosea sp. BK604* | 609 |
| A0A521LQC9 | Methyl-accepting transducer domain-containing protein | *Bosea sp. (in: a-proteobacteria)* | 610 |
| A0A542B8V0 | Methyl-accepting transducer domain-containing protein | *Bosea sp. AK1* | 610 |
| A0A5C1AZE6 | Methyl-accepting transducer domain-containing protein | *Bosea sp. F3-2* | 608 |
| A0A653N1K2 | Methyl-accepting transducer domain-containing protein | *Bosea sp. 62* | 620 |
| A0A6F8V2D2 | Methyl-accepting transducer domain-containing protein | *Bosea sp. ANAM02* | 610 |
| A0A840BX31 | Chemotaxis protein | *Chelatococcus caeni* | 620 |
| A0A841KDD1 | Methyl-accepting chemotaxis sensory transducer with Pas/Pac sensor | *Chelatococcus composti* | 620 |
| A0A922Y3U9 | Methyl-accepting transducer domain-containing protein | *Methylobacterium sp.* | 611 |
| A0A922YZ18 | Methyl-accepting chemotaxis sensory transducer with Pas/Pac sensor | *Methylobacterium sp.* | 642 |
| A0A927EAW8 | Methyl-accepting transducer domain-containing protein | *Bosea spartocytisi* | 607 |
| A0A950HSZ1 | PAS domain S-box protein | *Methylobacteriaceae bacterium* | 613 |
| A0A9D8LXK7 | Methyl-accepting transducer domain-containing protein | *Bosea sp. (in: a-proteobacteria)* | 609 |
| A0A9D8Q8D5 | Methyl-accepting transducer domain-containing protein | *Bosea sp. (in: a-proteobacteria)* | 609 |
| A0A9E7ZUZ9 | Methyl-accepting transducer domain-containing protein | *Bosea sp. NBC_00550* | 607 |
| A0A9E8CS09 | Methyl-accepting transducer domain-containing protein | *Bosea sp. NBC_00436* | 610 |
| A0A9W4W669 | Methyl-accepting chemotaxis protein | *Hyphomicrobiales bacterium* | 608 |
| A0A0A0I8T3 | Methyl-accepting transducer domain-containing protein | *Clostridium novyi A str. 4552* | 534 |
| A0A0A0IFN5 | Methyl-accepting transducer domain-containing protein | *Clostridium botulinum C/D str. DC5* | 535 |
| A0A0A0IR06 | Methyl-accepting transducer domain-containing protein | *Clostridium novyi A str. 4570* | 535 |
| A0A0E3K120 | Methyl-accepting transducer domain-containing protein | *Clostridium scatologenes* | 506 |
| A0A0Q6KRQ8 | Methyl-accepting transducer domain-containing protein | *Bosea sp. Leaf344* | 611 |

| | | | |
|---|---|---|---|
| A0A1B3NK79 | Methyl-accepting transducer domain-containing protein | *Bosea sp. RAC05* | 610 |
| A0A1H7RG25 | Methyl-accepting transducer domain-containing protein | *Bosea lupini* | 611 |
| A0A258A0I4 | Methyl-accepting transducer domain-containing protein | *Bosea sp. 12-68-7* | 610 |
| A0A258CVI6 | Methyl-accepting transducer domain-containing protein | *Bosea sp. 32-68-6* | 610 |
| A0A2G6QKH2 | Methyl-accepting transducer domain-containing protein | *Treponema sp.* | 570 |
| A0A2U8DYN4 | Methyl-accepting transducer domain-containing protein | *Clostridium drakei* | 535 |
| A0A6P1Y380 | Methyl-accepting transducer domain-containing protein | *Treponema vincentii* | 566 |
| A0A839K500 | Methyl-accepting transducer domain-containing protein | *Variimorphobacter saccharofermentans* | 540 |
| A0A8I1QZ84 | Methyl-accepting transducer domain-containing protein | *Bosea sp. (in: a-proteobacteria)* | 607 |
| A0A916PHV9 | Methyl-accepting transducer domain-containing protein | *Clostridium botulinum C str. Eklund* | 535 |
| A0A922YY04 | Methyl-accepting transducer domain-containing protein | *Methylobacterium sp.* | 610 |
| A0A9Q1UXK4 | Methyl-accepting transducer domain-containing protein | *Clostridium botulinum* | 535 |
| A0A9Q3VEX0 | Methyl-accepting transducer domain-containing protein | *Clostridium botulinum C/D* | 535 |
| A0Q1C1 | Methyl-accepting transducer domain-containing protein | *Clostridium novyi (strain NT)* | 535 |
| C6PMW7 | Methyl-accepting transducer domain-containing protein | *Clostridium carboxidivorans P7* | 536 |
| E7NQZ5 | Methyl-accepting transducer domain-containing protein | *Treponema phagedenis F0421* | 565 |

**Supplementary Table 7** 5TMR-sensor histidine kinases fused with a methyl-accepting transducer domain-containing protein (MCP). Receptors whose domain architecture includes also a PAS_3 domain are shaded in orange, while in green are SHKs formed exclusively from the 5TMR fold and the MCP domain. The protein list was obtained from https://www.ebi.ac.uk/interpro/entry/pfam/PF07694/domain_architecture/.
